# Genetic variation at 27 y-strs in four regions of bahrain

**DOI:** 10.1101/787341

**Authors:** Noora R. Al-Snan, Safia A. Messaoudi, Yahya M. Khubrani, Jon H. Wetton, Mark A. Jobling, Moiz Bakhiet

**Affiliations:** Department of Molecular Medicine, College of Medical and Medicine Sciences, Arabian Gulf University, Kingdom of Bahrain; Forensic Science Laboratory, Directorate of Forensic Science, General Directorate of Criminal Investigation and Forensic Science, Ministry of Interior, Kingdom of Bahrain; Forensic Sciences Department, College of Criminal Justice, Naif Arab University for Security Sciences, Riyadh, Saudi Arabia; Department of Genetics & Genome Biology, University of Leicester, University Road, Leicester, United Kingdom; Forensic Genetics Laboratory, General Administration of Criminal Evidence, Public Security, Ministry of Interior, Saudi Arabia

**Keywords:** Bahraini population, Y-STRs, Haplogroup, Population structure, Haplotype

## Abstract

Bahrain location in the Arabian Gulf contributed to the diversity of its indigenous population descended from Christian Arabs, Persians (Zoroastrians), Jews, and Aramaic-speaking agriculturalists. The aim of this study was to examine population substructure within Bahrain using the 27 Y-STRs (short tandem repeats) in the Yfiler Plus kit and to characterize the haplotypes of 562 unrelated Bahraini nationals, sub-divided into the four geographical regions - North, Capital, South and Muharraq. Yfiler Plus provided a significant improvement over the earlier 17-locus Yfiler kit in discrimination capacity, increasing it from 77% to 87.5%, but this value differed widely between regions from 98.4% in Muharraq to 75.2% in the Northern region, an unusually low value possibly as a consequence of the very rapid expansion in population size in the last 80 years. Clusters of closely related male lineages were seen, with only 79.4% of donors displaying unique haplotypes and 59% of instances of shared haplotypes occurring within, rather than between, regions. Haplogroup prediction indicated diverse origins of the population with a predominance of haplogroups J2 and J1, both typical of the Arabian Peninsula, but also haplogroups such as B2 and E1b1a originating in Africa and H, L and R2 indicative of migration from the east. Haplogroup frequencies differed significantly between regions with J2 significantly more common in the Northern region compared with the Southern possibly as a result of differential settlement with Baharna (descended from populations in which J2 predominates) and Arabs (both indigenous and migrant Huwala who are expected to have a higher frequency of J1). Our study illustrated the importance of encompassing geographical and social variation when constructing population databases and the need for highly discriminating multiplexes where rapid expansions have occurred within tightly bounded populations.

## Introduction

The Kingdom of Bahrain is the smallest archipelago in the Arabian Peninsula totaling just 765 square kilometers, located northwest of the State of Qatar, and east of the Kingdom of Saudi Arabia; further to the north and east lies the Islamic Republic of Iran **(1)**. Bahrain’s population of ∼ 1.6 million in 2019, less than half of whom are Bahraini citizens **(2)**, reside primarily on the main islands of Bahrain, Muharraq, Umm al-Naasan and Sitra **(3)**. Its coastal location in the Arabian Gulf, fertile land and abundance of fresh water have attracted many migrants resulting in an ethnically diverse society with origins both in the Arabian Peninsula and further afield such as Iran and India **(4)**. Bahrain’s pre-Islamic population consisted of Christian Arabs, Persians (Zoroastrians), Jews, and Aramaic-speaking agriculturalists **(5)** and now comprises four main ethnic groups: Arabs, Baharna (the putative indigenous peoples) and Persians (Huwala and Ajam) which are distributed unevenly between the four Governorates (Capital, Muharraq, Northern and Southern) **(6, 7)**. Of these, the Arabs have traditionally lived in areas such as Zallaq, Hawar Islands, Riffa (all in the Southern Governorate) and Muharraq whilst the Ajam, who are ethnic Persians **(8)**, and Baharna, the Arabized descendants of the pre-Islamic population, form large communities in the Capital Governorate, Muharraq and in some parts of the Northern Governorate. Muharraq and Riffa also have significant numbers of Huwala, descendants of migrant Arabs many of whom had journeyed from the Arabian Peninsula to Iran around the 18th or 19th century, largely returning between 1850 and 1900, as well as Bahrainis of Balochi descent and others of East African origin **(5)**. This geographical and social organization might be expected to have an effect on patterns of genetic diversity, particularly at loci on the male-specific region on the Y chromosome (MSY), which tends to show pronounced geographic variation **(9-11)**. To date genetic studies of the Bahraini population have been limited and little has been done to characterize population structure within the Kingdom.

Here we use Yfiler Plus to characterize Y-STR (short tandem repeat) haplotypes in 562 unrelated male Bahraini citizens sub-divided by geographical region, the same samples have been previously typed with autosomal STRs **(12)**. The 6-dye Yfiler Plus PCR Amplification kit detects 27 Y-STR loci; DYS19, DYS385a/b, DYS389I/II, DYS390, DYS391, DYS392, DYS393, DYS437, DYS438, DYS439, DYS448, DYS456, DYS458, DYS460, DYS481, DYS533, DYS635 (Y-GATA-C4), Y-GATA-H4, DYF387S1a/b, DYS449, DYS518, DYS570, DYS576 and DYS627, with the last seven loci being rapidly mutating Y-STRs (RM Y-STRs). These RM-YSTRs, which were not detected by the original Yfiler multiplex, are particularly useful for distinguishing between closely related males **(13-15)** and increasing the likelihood of discrimination in populations that have grown very rapidly often from small numbers of patrilinealy related tribal groups **(16)**.

## Materials and methods

### Sample Collection

Blood spots were collected on Nucleic-Cards (Copan, Italy) from 562 unrelated Bahraini males whose ancestry to the level of paternal grandfather was assigned to one of four administrative subdivisions of the country (Capital, Muharraq, Northern and Southern Governorates). Donors ranging in age from 20 to 55 years were recruited through social media channels such as Twitter and Instagram and invited to the General Directorate of Criminal Investigation and Forensic Science – Kingdom of Bahrain to submit blood samples. Ethical review for the study was provided by the Research and Research Ethics Committee (RREC) (E007-PI-10/17) in the Arabian Gulf University and informed consent was provided by all participants along with details sufficient to exclude individuals sharing ancestry closer than paternal great grandfather.

### DNA amplification and fragment detection

DNA was obtained from 1.2-mm diameter discs punched from blood-spots using the easyPunch STARlet system (Hamilton). A total of 27 Y-STRs were directly amplified with the Yfiler Plus Amplification Kit using 28 cycles according to manufacturer’s recommendation on a Veriti 96- Well Thermal Cycler (Thermo Fisher Scientific). The PCR products (1 µl) were separated by capillary electrophoresis in an ABI 3500xl Genetic Analyzer along with LIZ600 size standard v2 in a Hi-Di Formamide master mix. GeneMapper ID-X Software v1.4 was used for genotype assignment using the allelic ladders provided with the Yfiler Plus kit following ISFG recommendations **(17)** Samples displaying non-standard patterns, off-ladder and microvariant alleles were repeated. The resulting profiles were submitted to the Y-chromosomal Haplotype Reference Database **(18)** under the following accession numbers: Capital Governorate YA004556, Northern Governorate YA004557, Southern Governorate YA004558 and Muharraq Governorate YA004559.

### Statistical analysis

#### Forensic and population genetic parameters

Haplotype information (number of haplotypes, number of unique haplotypes, discriminatory capacity and haplotype diversity) was calculated using GenAlEx v6.503 in which all DNA samples were compared using the haplotype matching function available in the software **(19)**.

Allele and haplotype frequencies were estimated using STRAF **(20)**. Genetic diversity (GD) was calculated according to Nei **(21)**, haplotype match probability (HMP) was calculated as the sum of the squared haplotype frequencies, and the discriminatory capacity (DC) was calculated as the ratio between the number of different haplotypes and the total number of haplotypes. Haplotype diversities (HD) were calculated as one minus the HMP multiplied by the number of haplotypes, divided by the number of haplotypes minus one. Comparison of the forensic and the genetic parameters of the AmpFlSTR Yfiler and the Yfiler plus loci was performed with the ySTRmanager online tool **(22)**.

Genetic distances between populations were evaluated with R_st_ and visualized through multi-dimensional scaling (MDS) plots using comparative population data and the calculation tool within the online YHRD **(23)**. Population differentiation tests based on predicted haplogroup frequencies were carried out within Arlequin **(24)**.

For all statistical analyses, the alleles of the DYS389 locus were converted to the DYS389p & DYS389q nomenclature by subtracting the repeat number of DYS389I region from that of DYS389II so that its diversity was not considered twice.

### Haplogroup prediction

The NevGen Y-DNA Haplogroup Predictor **(25)**, based on a previously-implemented Bayesian approach **(26)**, was used to derive likely Y-SNP haplogroups from Y-STR haplotypes **(25)**. The software bases the prediction upon length variation at the 23 loci in the Promega PowerPlex Y23 multiplex, so DYS549 and DYS643 (which are not amplified by Yfiler Plus) were coded as missing data. Whilst the Predictor allocates haplotypes to one of 484 sub-branches of the Y haplogroup tree, we have restricted our assignments to major haplogroups which we have previously shown can be accurately predicted in Arabian Peninsula populations **(16, 27)**.

### Median-joining networks

Median-joining networks of haplotypes were constructed using the program Network v. 5.0.1.1 **(28)** with weightings based on the inverse variance of repeat length at each locus in order to reduce reticulation. The duplicated loci DYS385a/b and DYF387S1a/b were excluded from the network construction as it is not possible to associate particular alleles to specific copies.

## Results

### Haplogroup prediction & median–joining networks

A median-joining network based solely on allele length differences between the haplotypes of the 562 Bahraini males’ displays several distinct branches as shown in **Figs (1-3)**. An almost perfect correspondence was noted between these and the haplogroup predictions. Haplogroup prediction suggests that haplogroup J2 is the most common in the Bahraini population encompassing 27.6% of men, followed by J1 (23.0%), E1b1b (8.9%), E1b1a (8.6%) and R1a (8.4%), with other predicted haplogroups (G, T, L, R1b, Q, R2, B2, E2 and H) occurring at progressively lower frequencies. The profiles of 20 men were assigned low haplogroup fitness (<25) and a high probability of haplogroup misassignment (>97.5%) because there were no closely matching haplotypes of known haplogroup within the NevGen prediction model; these were therefore designated as unpredicted (UP).

**Figure 1.**
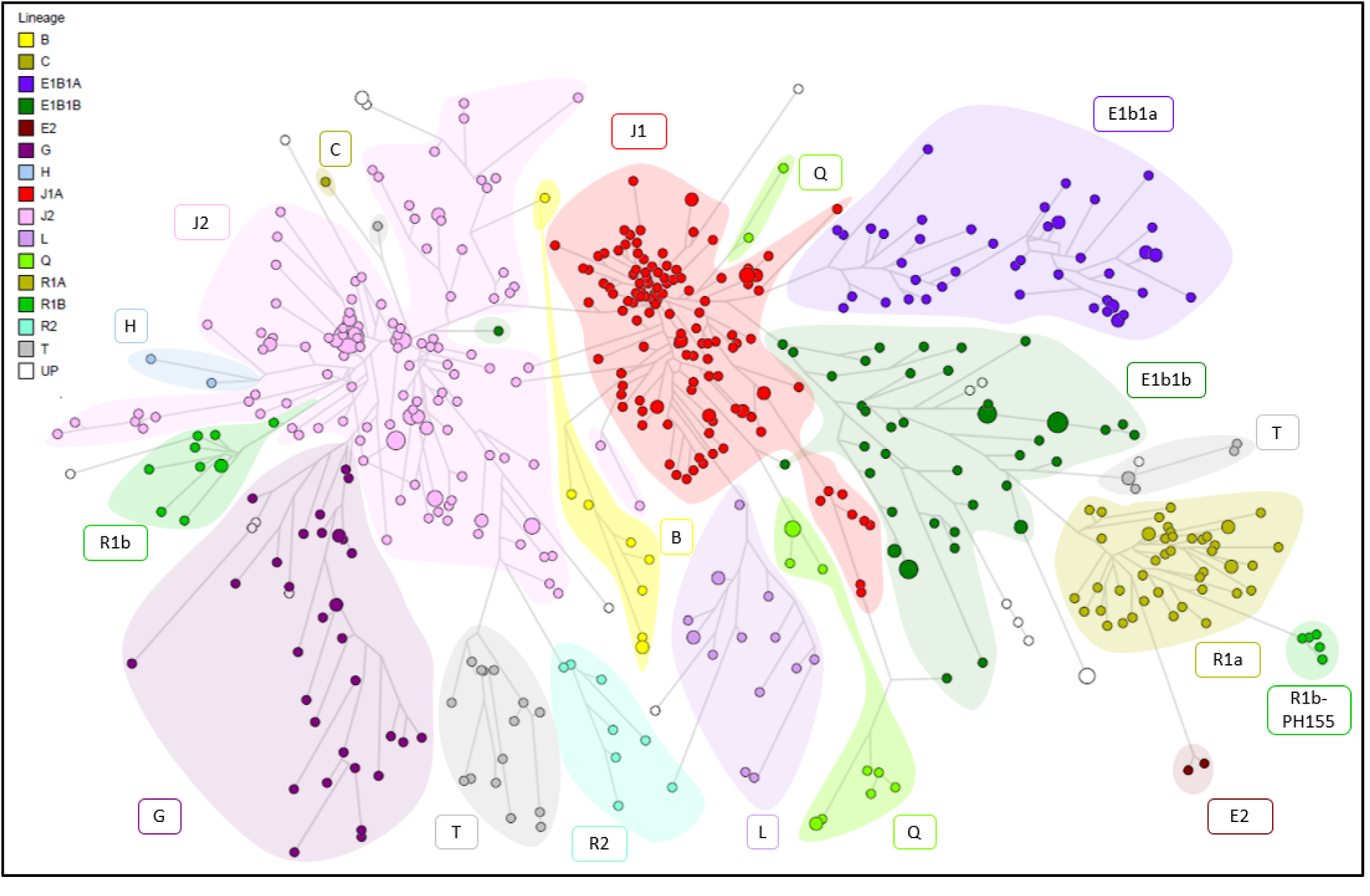
Distribution of Y-STR haplogroups in the Bahraini population.

**Figure 2.**
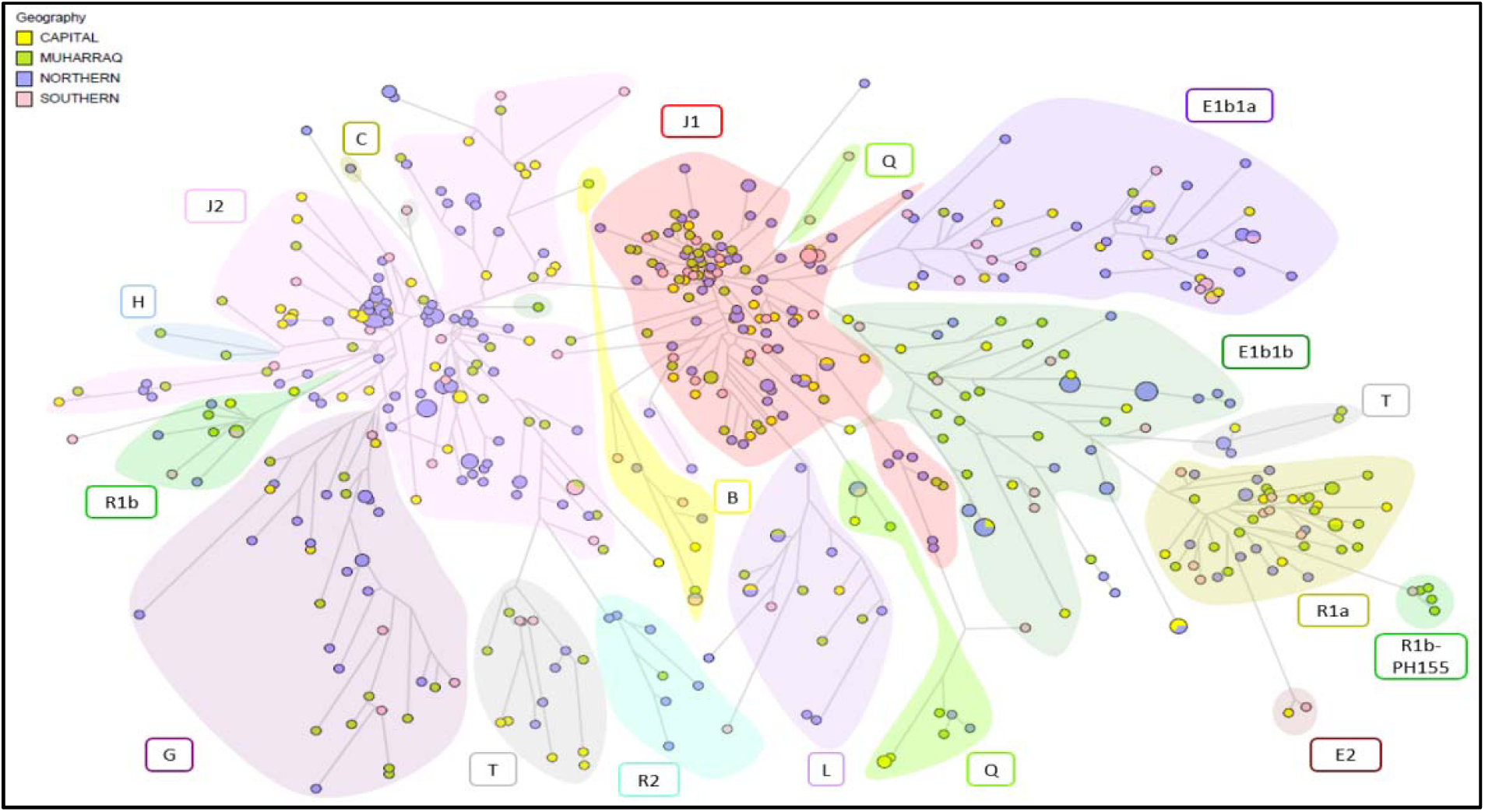
Distribution of Y-STR haplogroups within the regions of Bahrain.

**Figure 3.**
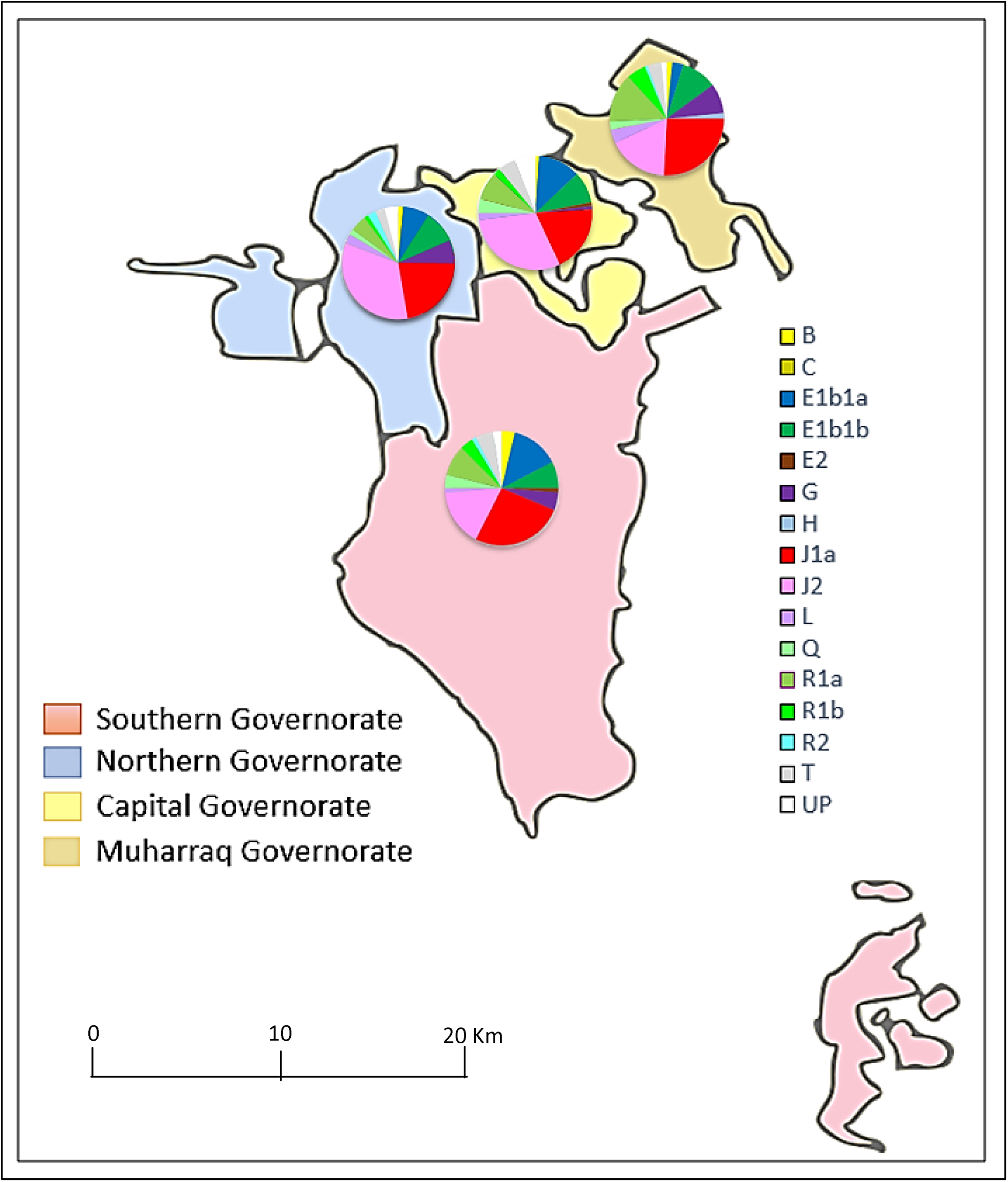
Distribution of YSTR haplogroups between different regions.

Haplogroup predictions allow comparison with SNP-defined lineages that are known from published studies to differ significantly in frequency between countries. **Table 1** demonstrates that haplogroup frequency differences also exist between governorates. The ratio of J1 to J2 shows clear differences between the governorates (Fisher’s Exact Test P= 0.012). J1 is most frequent in the Southern Governorate (27%) where the highest proportion of Arabs live, and in the Muharraq Governorate (27%) where many migrant Huwala Arabs resettled, and it declines to its lowest frequency in the Northern and Capital Governorates (21% and 19%). In contrast the Northern and Capital Governorates where the Baharna and Ajam are most common show higher frequencies of haplogroup J2 (34% and 31%) than in Muharraq and the Southern Governorate (both 17%).

**Table 1.** Regional frequencies of clustered predicted Y-haplogroups for 562 Bahraini males.

|  |  | Predicted haplogroup |  |  |  |  |  |  |  |  |  |  |  |  |  |  |  |
| --- | --- | --- | --- | --- | --- | --- | --- | --- | --- | --- | --- | --- | --- | --- | --- | --- | --- |
| Region | N | B | C | E1b1a | E1b1b | E2 | G | H | J1a | J2 | L | Q | R1a | R1b | R2 | T | UP |
| Capital | 100 | 1 |  | 12 | 9 | 1 | 1 |  | 19 | 30 | 2 | 4 | 8 | 2 |  | 5 | 6 |
| Muharraq | 128 | 2 |  | 4 | 13 |  | 11 | 2 | 33 | 22 | 5 | 3 | 18 | 7 | 1 | 5 | 2 |
| Northern | 254 | 3 | 1 | 19 | 24 |  | 17 |  | 56 | 85 | 7 | 4 | 12 | 3 | 6 | 7 | 10 |
| Southern | 80 | 3 |  | 11 | 6 | 1 | 4 |  | 21 | 13 | 1 | 3 | 7 | 3 | 1 | 4 | 2 |
| Bahrain | 562 | 9 | 1 | 46 | 52 | 2 | 33 | 2 | 129 | 150 | 15 | 14 | 45 | 15 | 8 | 21 | 20 |

**N:** number of individuals; **UP:** unpredicted haplogroup

Pairwise F_ST_ based on predicted haplogroup frequency shows the greatest overall differences are between Muharraq and Northern Governorates (F_ST_ =0.01906, P<0.0001) and Muharraq and Capital Governorates (F_ST_ =0.01586, P<0.0001) with other comparisons not reaching formal significance after Bonferroni correction **(Table 2)**. Similarly Fisher’s Exact Tests of population differentiation highlighted the same pairs as being significant with the comparison between the Northern and Southern Governorates once again just failing to reach significance after Bonferroni correction. A similar pattern was seen when considering R_ST_ based on the haplotypes themselves but in this case whilst Muharraq and Northern Governorates (R_ST_ =0.01179, P<0.0001) remained highly significant the other comparisons were not following Bonferroni correction **(Table 3)**.

**Table 2.** FST values for pairwise comparisons of predicted haplogroup frequency.

|  | Capital | Northern | Muharraq | Southern |
| --- | --- | --- | --- | --- |
| Capital | 0 | 0.3603+-0.058 | **<0.00001** | 0.1441+-0.034 |
| Northern | 0 | 0 | **<0.00001** | 0.0180+-0.012 |
| Muharraq | 0.01586 | 0.01906 | 0 | 0.3784+-0.030 |
| Southern | 0.00555 | 0.01599 | 0.00051 | 0 |

**Table 3.** RST values for pairwise comparisons of haplotypes between populations.

|  | Capital | Northern | Muharraq | Southern |
| --- | --- | --- | --- | --- |
| Capital | 0 | 0.0811+-0.037 | 0.0451+-0.020 | 0.3333+-0.051 |
| Northern | 0.00404 | 0 | **<0.00001** | 0.0180+-0.012 |
| Muharraq | 0.00736 | 0.01179 | 0 | 0.2162+-0.043 |
| Southern | 0.00211 | 0.00894 | 0.00182 | 0 |

F_ST_ values for pairwise comparisons of predicted haplogroup frequency, shown below the diagonal and probabilities shown above, those which were significant after Bonferroni correction are shown in bold.

R_ST_ values for pairwise comparisons of haplotypes between populations shown below the diagonal and probabilities shown above, those which were significant after Bonferroni correction are shown in bold.

### Y-STR allele and haplotype diversity

Microvariant alleles aligning with the Yfiler Plus virtual allelic ladder were detected in 141 samples; 129 samples displayed .2 microvariants at DYS458 (14.2, 16.2, 17.2, 18.2, 19.2, 20.2, 21.2 and 22.2), which corresponded exactly with haplotypes predicted to belong to haplogroup J1 in concordance with the well-established association between intermediate length (.2) DYS458 alleles and this haplogroup **(16, 29)**. Four samples displayed .2 microvariants present in the DYF387S1 virtual ladder (39.2 and 41.2), and all were predicted to be haplogroup B2 which is often associated with .2 microvariants at this locus **(26)**. Similarly, a DYS481 25.1 microvariant in five men was wholly concordant with the NevGen prediction of the R1b-PH155 subclade, which is a globally rare basal R1b lineage previously recorded in several Bahrainis with similar haplotypes who have submitted their Y-STR and Y-SNP profiles to the YFull Tree database **(18, 30)**. The same allele (25.1) has also been recorded in a Han Chinese individual who was derived for M343 but ancestral for L389 which also implies a basal R1b lineage **(31)**.

Several off-ladder (OL) alleles were designated after comparison with the allelic ladder. Six OL alleles were detected at DYF387S1; a single example of 36.3 in a predicted G2a haplotype, 35.2 in two individuals sharing an identical R1a haplotype and a single instance of 44.2 and two occurrences of 42.2 in three different B2a haplotypes (these representing OL variants of the previously mentioned association of .2 alleles with the B2 haplogroup). Two occurrences of the OL DYS570 allele 14.3 were found in closely related J1a haplotypes differing by a single repeat at each of four loci. Three OL alleles were detected at DYS449 (two instances of 34.2 and a single 32.1) all within closely related predicted E1b1 lineages.

An unusual finding was an OL peak of 219 bp within the DYS19 size range in association with a standard DYS19 allele 15. This donor lacked a peak within the expected DYS448 size range raising the possibility that the OL peak is the result of an unusually large internal deletion producing a DYS448 allele 4. The donor was predicted to belong to haplogroup J2a which typically has alleles in the range of 19-23 repeats (309-334 bp). The common structure of alleles at this locus comprises two variable length (AGAGAT)n hexamer repeats separated by a 42-bp “non-variable” region of seven diverged hexamer repeats; the observed OL peak is appropriately sized to have resulted from the loss of between 15 and 19 hexamer repeats. A similar event has been reported previously by Budowle et al. **(32)**. As well as this “pseudo-null” DYS448 allele, a true null allele was detected at DYS439 in a donor of unpredicted haplogroup.

Duplications at DYF387S1 produced two instances of tri-allelic patterns similar to those reported in several populations **(26, 33-37)**. In one predicted J1a donor the alleles 36, 37 and 38 were all detected with equal strength implying an equal copy number as would be seen if just one DYF387S1 copy had been duplicated. The second donor who was predicted as E1b1b-V13 displayed alleles 35, 36 and 37, but the 35 peak was double the height of the others implying that there were four copies of the locus in this individual. The different haplogroup affiliations show that duplications in this region are both frequent and diverse events. Another possible mutational event associated with duplicated STR loci is gene conversion leading to the homogenization of the repeat number between two copies **(38)**. This process may explain why two predicted haplogroup B2 individuals share a haplotype with only a 37 repeat peak whilst the other seven predicted B2 haplotypes in this study all display one copy with an integer number of repeats differing by at least three repeats from an intermediate (.2) copy as is frequently observed in other B2 lineages **(26)**. It is likely there was a gene conversion back to the ancestral (integer) state earlier in the paternal lineage of these two men.

The complete haplotype list and diversity summary statistics are provided for the four subpopulations **(Table 4)**. As expected, the number of haplotypes was considerably increased by the ten additional loci compared with the earlier Yfiler multiplex; 492 distinct Yfiler Plus haplotypes were detected in the 562 samples. Of these, 446 haplotypes were observed once (90.7%), and there were 33 identical pairs, eight trios, three quartets, one quintet and a haplotype shared nine times (an ennead).

**Table 4.**
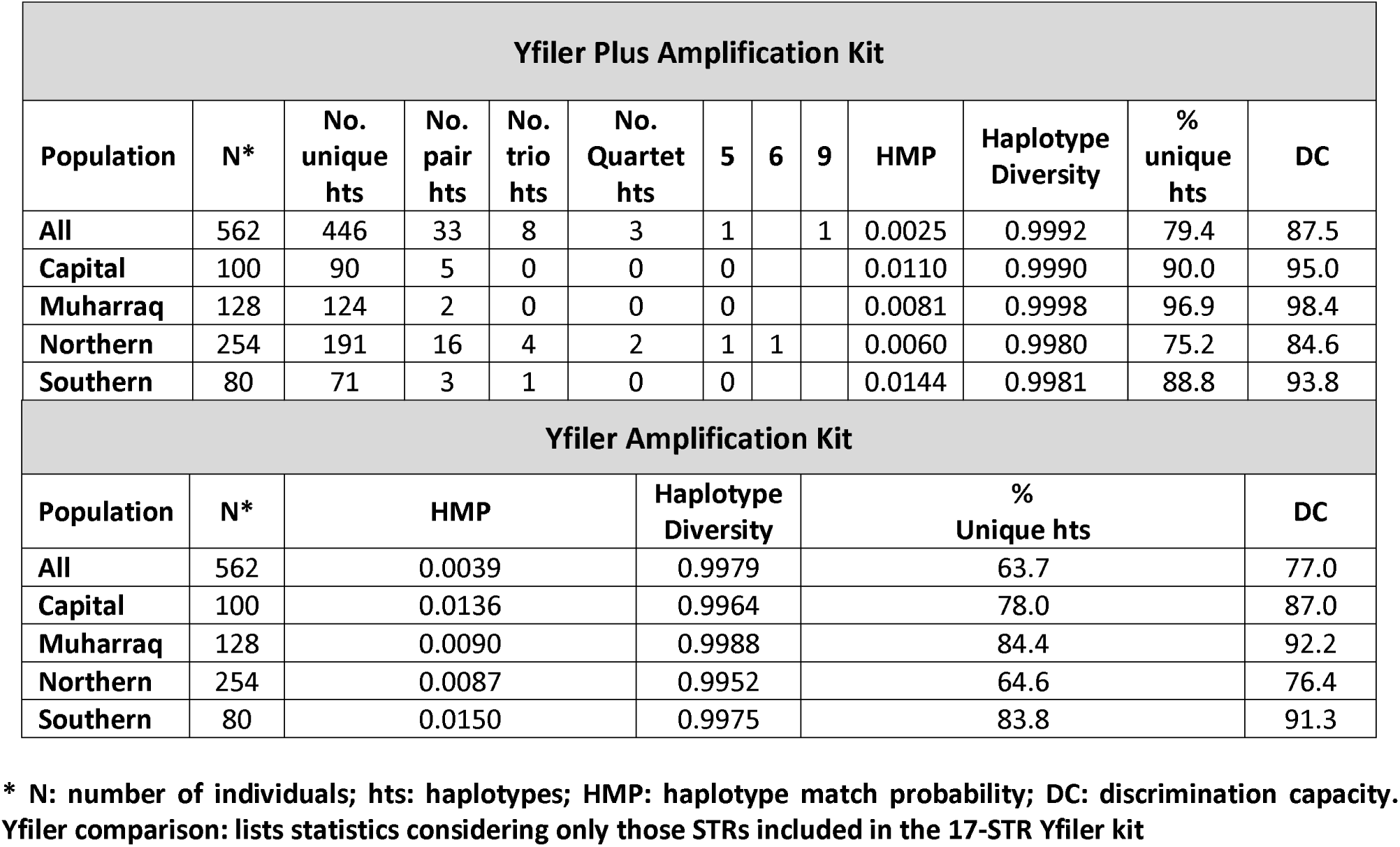
Diversity summary statistics for Y-STR haplotypes, considering region of recruitment.

| Yfiler Plus Amplification Kit |  |  |  |  |  |  |  |  |  |  |  |  |
| --- | --- | --- | --- | --- | --- | --- | --- | --- | --- | --- | --- | --- |
| Population | N* | No. unique hts | No. pair hts | No. trio hts | No. Quartet hts | 5 | 6 | 9 | HMP | Haplotype Diversity | % unique hts | DC |
| All | 562 | 446 | 33 | 8 | 3 | 1 |  | 1 | 0.0025 | 0.9992 | 79.4 | 87.5 |
| Capital | 100 | 90 | 5 | 0 | 0 | 0 |  |  | 0.0110 | 0.9990 | 90.0 | 95.0 |
| Muharraq | 128 | 124 | 2 | 0 | 0 | 0 |  |  | 0.0081 | 0.9998 | 96.9 | 98.4 |
| Northern | 254 | 191 | 16 | 4 | 2 | 1 | 1 |  | 0.0060 | 0.9980 | 75.2 | 84.6 |
| Southern | 80 | 71 | 3 | 1 | 0 | 0 |  |  | 0.0144 | 0.9981 | 88.8 | 93.8 |
| Yfiler Amplification Kit |  |  |  |  |  |  |  |  |  |  |  |  |
| Population | N* | HMP |  |  | Haplotype Diversity |  | % Unique hts |  |  | DC |  |  |
| All | 562 | 0.0039 |  |  | 0.9979 |  | 63.7 |  |  | 77.0 |  |  |
| Capital | 100 | 0.0136 |  |  | 0.9964 |  | 78.0 |  |  | 87.0 |  |  |
| Muharraq | 128 | 0.0090 |  |  | 0.9988 |  | 84.4 |  |  | 92.2 |  |  |
| Northern | 254 | 0.0087 |  |  | 0.9952 |  | 64.6 |  |  | 76.4 |  |  |
| Southern | 80 | 0.0150 |  |  | 0.9975 |  | 83.8 |  |  | 91.3 |  |  |

\* N: number of individuals; hts: haplotypes; HMP: haplotype match probability; DC: discrimination capacity. Yfiler comparison: lists statistics considering only those STRs included in the 17-STR Yfiler kit

Considering all the Yfiler Plus loci the proportion of donors within the national sample with unique haplotypes was 79.4%, declining to 63.7% when only the Yfiler loci were analyzed. The diverse Muharraq Governorate had the highest percentage of unique haplotypes (96.9% for Yfiler Plus, reducing to 84.4% for Yfiler), whilst the lowest percentage bearing unique haplotypes was seen in the Northern Governorate (75.2% reducing to 64.6% at the Yfiler loci). The majority of shared haplotypes were shared within rather than between regions (59% against an expectation of 31% if lineages were equally dispersed across the governorates).

This was naturally reflected in the discriminatory capacity (DC), where the national value for the Yfiler Plus loci was 87.5%, considerably higher than when using Yfiler (77.0%).

Regarding the subpopulations, Muharraq Governorate recorded the highest DC (98.4%) followed by Capital Governorate (95.0%), with Northern Governorate the lowest at 84.6%.

### MDS analysis

AMOVA was used to explore population differentiation with pairwise genetic distances (R_st_) visualized by multidimensional scaling (MDS), between the four Governorates of Bahrain and relevant neighboring population samples obtained from YHRD Release R61 **(Fig 4)**. Comparisons were performed using just the Yfiler loci as Yfiler Plus data were not available for Iran, the region from which a significant proportion of the Bahraini population had originated. Muharraq shows a greater degree of similarity to Iran whereas Southern, Northern and Capital populations were closer to Emirati and Saudi populations which perfectly correlated with autosomal results obtained by comparison with autosomal profiles from the same individuals **(12)**. Muharraq also fell closest to the only Bahraini data previously submitted to YHRD (YA004278, N=156 Yfiler profiles) although the regional origins of the donors to that study are unknown.

**Figure 4.**
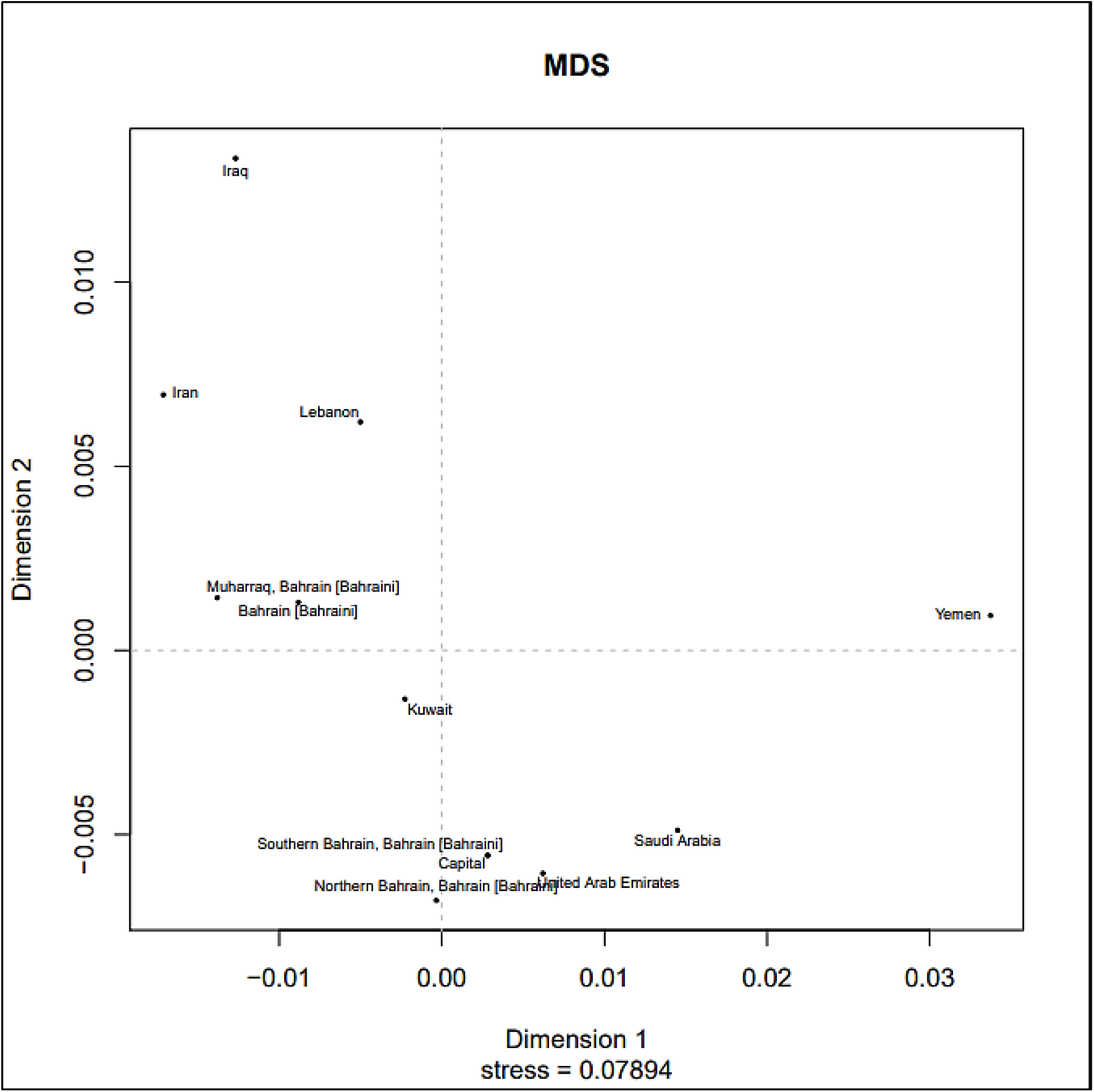
MDS plot between Bahraini and other populations.

## Discussion

While the addition of a number of RM Y-STRs greatly increased the discrimination capacity (DC) compared with the Yfiler multiplex previously implemented in Bahrain, a DC of 87.5% remains very low compared with most global populations such as Serbia 99.9% (39), Upper Austria and Salzburg 99.7% **(37)**, Mongolia 98.9% **(40)**, Italy 98.5% **(41)**, US Caucasians 98.5% **(42)**, Daur 96.55% **(40)** and Saudi Arabia 95.3% **(16)** although the highly bottle-necked Greenland population is lower at 79.0% **(43)**. There are many shared haplotypes in the Bahraini sample, despite precautions to exclude males sharing common patrilineal ancestry in the last three generations. This is likely the result of an extended period of rapid population expansion since the discovery of oil in the early 1930s. Increased prosperity since then has led to improvements in health care and a significantly reduced childhood mortality rate which, combined with a predominantly youthful population, has resulted in Bahrain having the tenth most rapidly growing population in the world **(44, 45)**.

Lying at the crossroads of Europe, Asia and Africa, the genetic landscape of Bahrain has been shaped by migrants from many other regions. Prior to the opening in 1986 of a 25-km causeway to Saudi Arabia, all international contact had been via maritime routes through the Arabian Gulf. Unlike its immediate neighbors Saudi Arabia and Qatar, where haplogroup J1 predominates (71% and 58% respectively **(26, 46)**, the frequency in Bahrain is just 23%, being highest in the largely desert Southern Governorate where ethnic Arabs are most common and declining in the more urban areas where the Persian Ajam and indigenous Baharna are most abundant. The diverse haplogroup composition hints at the variety of peoples who have left their mark on Bahrain with B2 & E1b1a originating in Africa and H, L and R2 indicative of migration from the east, while the R1b haplotypes may result from the period of Portuguese rule from 1521 to 1602. We observed haplotypes predicted to belong to both primary branches of R1b, namely R1b1a- L754 and R1b1b-PH155, whilst R1b1a is by far the commonest globally the five examples predicted to belong to the very scarce basal haplogroup R1b1b-PH155 all carried a distinctive 25.1 intermediate allele at DYS481, showing that distinctive rare variants can locally reach high frequencies and emphasizing the relevance of appropriate regional databases **(47)**.

## Conclusions

We have reported the genetic characterization of the 27 Y-STR loci in the Yfiler Plus multiplex among Bahraini nationals from each of the four governorates (Northern, Southern, Muharraq, and Capital). Significant differences exist between the regions in haplotype frequencies with the majority of shared haplotypes being seen within, rather than between, governorates.

This likely reflects differences in ethnic composition exacerbated by the extremely rapid population growth the country has experienced in the last 70 years, which has led to an unusually high proportion of closely related males. The much increased discrimination power offered by the inclusion of several RM Y-STRs in the Yfiler Plus kit is consequently a major advance in improving discrimination power in this country.

## Acknowledgments

We would like to thank the authorities in Bahrain Forensic Science Lab, Mr. Abdulaziz Mayoof Alrumaihi, Mr. Raed Ali Almaeeli and Mr. Mohammed A. Ghayyath for allowing us to utilize the Bahrain Forensic Science Laboratory.

Also, many thanks to Latifa Ahmed and Sabah Nazir for their technical support. Lastly, many thanks to Dr. Mohammad A. Alenizi, Mr. Yasser Alaraibi for their valuable feedbacks and continuous support. This research did not receive any specific grant from funding agencies in the public, commercial, or not-for-profit sectors.

